# Dual-Stream Compression of High Bit-Depth Medical Images with Application to DNA Storage

**DOI:** 10.64898/2026.05.17.724501

**Authors:** Hongyi Su, Weikun Fan, Jialin Peng, Yanju Zhang

**Author notes:** Corresponding author: Yanju Zhang.

## Abstract

High bit-depth medical images preserve subtle intensity variations that are important for quantitative analysis and clinical interpretation, but their large dynamic range poses challenges for efficient compression. We propose a bit-plane-aware dual-stream compression framework for 16-bit medical images by separately modeling the most significant bit (MSB) and least significant bit (LSB) components. The MSB structural stream is encoded using JPEG coding with a Duplicate Segment Skipping (DSS) strategy to exploit spatial and segment-level redundancy, while the LSB detail stream is compressed using learned image compression to represent residual variations and fine-grained details. Experiments on four MRI and CT datasets show that the proposed method consistently outperforms representative traditional and learning-based codecs, achieving the lowest bit rate across all datasets. Meanwhile, it preserves high reconstruction fidelity. As a downstream application, we further demonstrate that the compressed bitstreams can be effectively integrated with DNA encoding and converted into sequences with favorable biochemical properties.

## I. Introduction

The exponential growth of clinical imaging data poses an unprecedented challenge to healthcare storage and long-term archiving systems [1]. Medical images are commonly stored with high bit-depth, such as 16-bit DICOM images, to preserve subtle intensity variations that are important for quantitative analysis and clinical interpretation [2]. Compared with natural 8-bit images, high bit-depth medical images have a much larger dynamic range and more complex intensity distributions, making efficient compression more challenging [3]. Therefore, developing effective compression methods for 16-bit medical images is essential for reducing storage costs while preserving diagnostically relevant information.

Existing image compression methods can be broadly divided into traditional codecs and learning-based approaches. Traditional codecs provide stable reconstruction quality and are easy to deploy [4], [5], while learning-based compression methods have achieved promising performance by using neural transforms and learned entropy models [6], [7]. However, many of them are primarily developed for natural images or general compression scenarios, and may not fully exploit the specific information distribution and diagnostic requirements of 16-bit medical images. These limitations highlight the need for a compression framework tailored to high bit-depth medical data.

A key observation in high bit-depth medical images is that different bit planes carry different types of information [8]. The most significant bits (MSB) mainly represent dominant anatomical structures and global intensity patterns, whereas the least significant bits (LSB) contain fine residual variations, local textures, and low-amplitude details. This motivates us to decompose 16-bit medical images into MSB and LSB components and compress them with different strategies.

Based on this observation, we propose a bit-plane-aware dual-stream compression framework for 16-bit medical images. The MSB structural stream is compressed using JPEG coding combined with a Duplicate Segment Skipping (DSS) strategy, which removes redundant restart-marker-delimited segments while preserving major anatomical structures. The LSB detail stream is compressed using a Learned Image Compression (LIC) model to obtain a compact representation of residual details. In addition to conventional image compression, we further evaluate the proposed framework as an image compression stage before DNA encoding, where reducing the compressed bitstream directly decreases the number of synthesized nucleotides and storage cost.

The main contributions of this work are summarized as follows:

- We propose a bit-plane-aware dual-stream compression framework for 16-bit medical images. The MSB structural stream is encoded using JPEG coding with DSS, while the LSB detail stream is compressed using learned image compression. By separately modeling high-order structural information and low-order residual details, the proposed method achieves a favorable balance between compression efficiency and reconstruction fidelity.
- We comprehensively evaluate the proposed method on multiple medical image datasets covering different modalities and anatomical regions. Experimental results demonstrate that the proposed method achieves superior compression performance while preserving pixel-level fidelity.
- We further investigate the application of the proposed compression framework to DNA-based medical image storage. The results show that the proposed method can substantially reduce the number of required nucleotides and improve coding density, demonstrating its potential as an effective preprocessing step before DNA encoding.

## II. Related work

Traditional medical image compression methods mainly rely on hand-crafted transforms, predictive coding, and entropy coding. Lossless or near-lossless codecs such as JPEG-LS [4], JPEG-2000 [5], PNG [9], JPEG-XL [10], and JP3D [11] have been widely used or investigated for medical image archiving due to their mature implementations, standardization, and reliable reconstruction. These methods are effective in exploiting spatial redundancy and preserving pixel-level fidelity. However, their compression efficiency is often limited when directly applied to high bit-depth medical images with complex intensity distributions, especially when large-scale storage or transmission is required.

Learning-based image compression methods have shown strong potential by replacing manually designed coding modules with learned probability models, transform-domain representations, and neural entropy models. Representative methods include L3C [12], LC-FDNet [13], aiWave [14], BCM-Net [15], and BD-LVIC [16]. Specifically, L3C introduces a learned hierarchical probabilistic model for full-resolution lossless image compression, while LC-FDNet improves learned lossless compression through frequency decomposition and coarse-to-fine modeling. For volumetric medical images, aiWave learns a 3-D affine wavelet-like transform to better capture spatial correlations, and BCM-Net improves lossless 3D medical image compression by modeling residuals with bilateral contextual information. BD-LVIC is most closely related to our work, as it adopts a bit-division strategy for high bit-depth medical volumes, using traditional codecs for the most significant bit-volume and learned entropy modeling for the least significant bit-volume.

However, most of them are designed for either general lossless image compression or strictly lossless volumetric medical image compression. In contrast, our work explores a high-fidelity compression setting for high bit-depth medical image slices, aiming to improve compression efficiency while preserving image structures that are relevant to medical interpretation. Rather than enforcing exact pixel-wise reconstruction, the proposed framework adopts a hybrid MSB/LSB coding design to separately compress high-order structural information and low-order residual details. The reconstruction quality is evaluated using reconstruction metrics, together with compression efficiency metrics.

### III. Method

### A. Datasets and Preprocessing

In this study, we evaluate the proposed compression framework on four high bit-depth medical image datasets, including TRABIT [17], Chaos-CT [18], Covid-CT [19], and Heart-MRI [2]. These datasets cover different imaging modalities, including MRI and CT, as well as different anatomical regions such as the brain, abdomen, lung, and heart. The details of the datasets used in this study are summarized in Table I. The resolution is reported as width × height, and the number in parentheses denotes the number of slices used in the experiments.

**TABLE I.** Summary of the medical image datasets used in this study.

| Dataset | Modality | Position | Resolution (Slices) | Bit Depth | Signed |
| --- | --- | --- | --- | --- | --- |
| TRABIT | MRI | Brain | 208 × 176 (176) | 16-bit | × |
| Chaos-CT | CT | Abdomen | 512 × 512 (78) | 16-bit | × |
| Covid-CT | CT | Lung | 512 × 512 (45) | 16-bit | ✓ |
| Heart-MRI | MRI | Heart | 320 × 320 (100) | 16-bit | × |

For signed high bit-depth images, such as Covid-CT, an intensity offset is applied before bit-plane decomposition to map the pixel values into a non-negative range. The inverse offset is applied after reconstruction to recover the original intensity domain. This preprocessing ensures that the proposed bit-plane decomposition can be consistently applied to both signed and unsigned medical images.

In order to reduce the processing difficulty of high bit-depth medical images *I*, we divided the image into two low bit-depth components, narrowing the range of pixel values [20]. These two components were represented as the MSB component *I*_*M*_ :

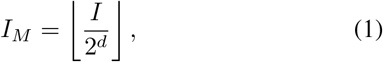

and the LSB component *I*_*L*_:

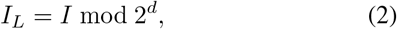

where *d* is a predefined integer, such as 4, 6 or 8.

### B. Compression of MSB Component

As illustrated in Fig. 1, the MSB component usually contains the dominant structural information of the image and exhibits piecewise-smooth characteristics. Compared with the LSB component, it has stronger spatial redundancy and is therefore more suitable for lightweight traditional coding. To preserve the major anatomical structures while maintaining low computational complexity, we employ JPEG compression [21] for the MSB stream. To further improve coding efficiency, we introduce a Duplicate Segment Skipping (DSS) strategy to remove redundant compressed segments in structurally uniform regions.

**Fig. 1.**
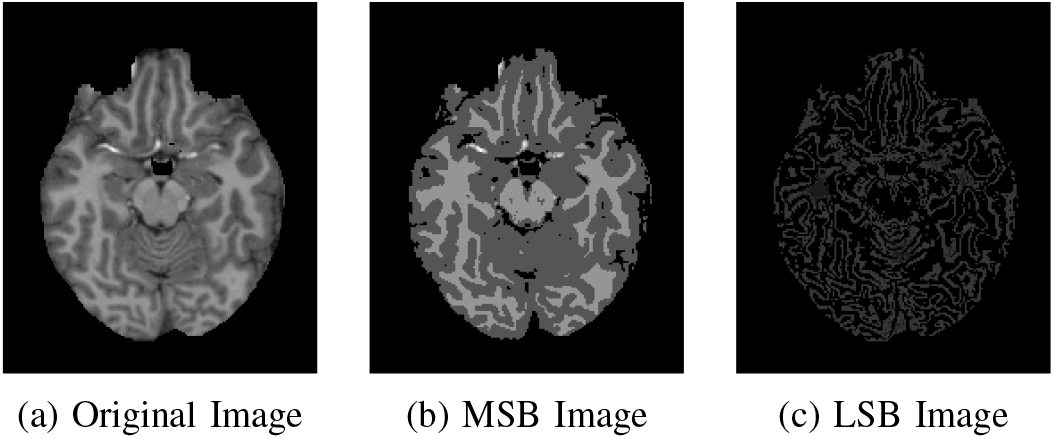
Example of an original high bit-depth medical image, and its corresponding MSB and LSB components.

As shown in Fig. 2, the MSB component is first encoded into a JPEG compressed scan stream with restart (RST) markers, which divide the entropy-coded stream into independently decodable segments. DSS is then applied to these restart-marker-delimited segments. Specifically, consecutive segments with identical compressed representations are regarded as duplicate segments. For each duplicate run, only the first segment and its original index are retained, while the subsequent repeated segments are skipped. During decoding, the recorded indices are used to identify the skipped positions, and the missing segments are restored by propagating the preceding valid JPEG segment. The complete JPEG stream is then reassembled and decoded to reconstruct the MSB component.

**Fig. 2.**
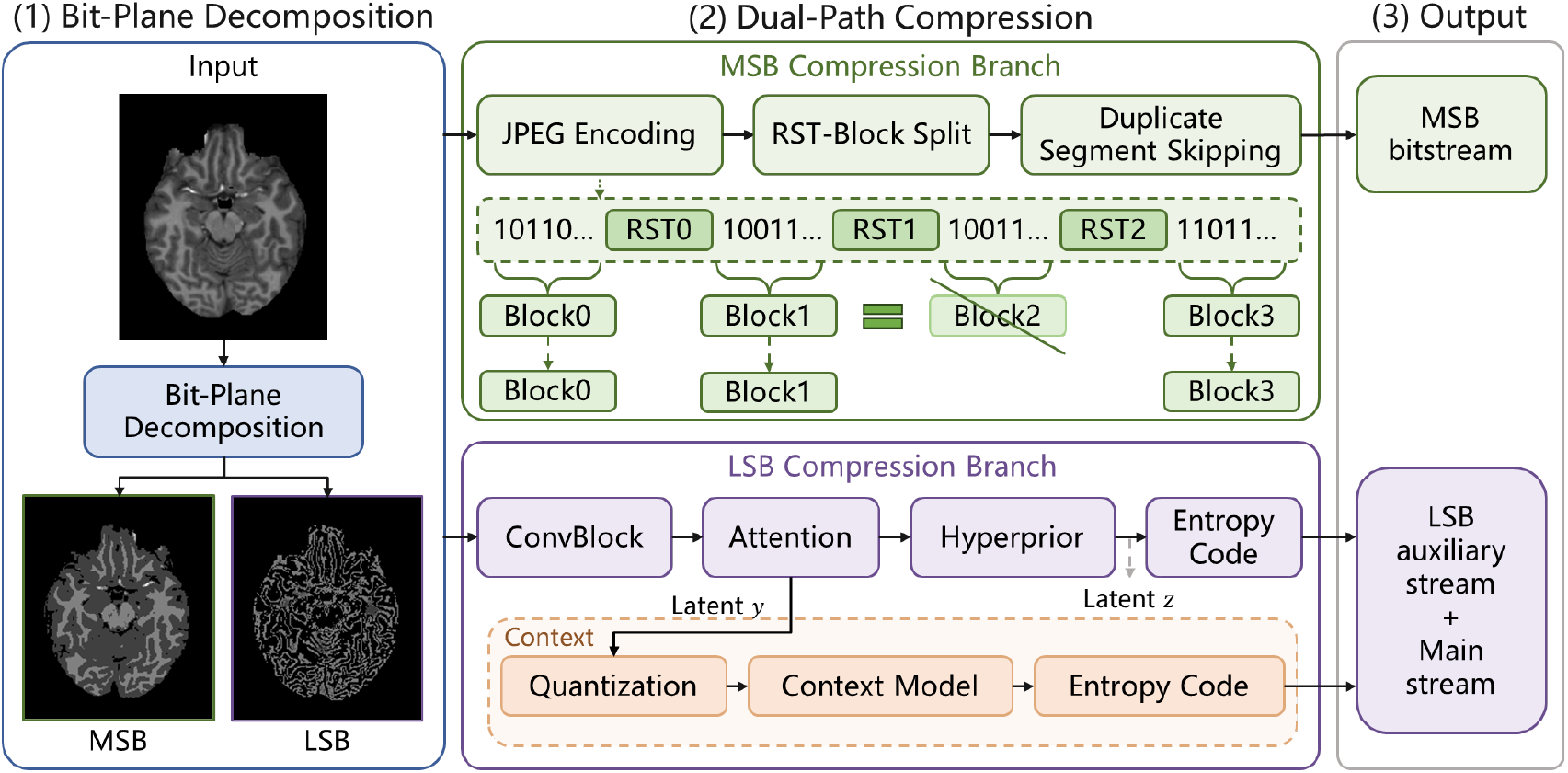
Overview of proposed method.

By operating on duplicate compressed segments without modifying the JPEG coding process itself, DSS further reduces the amount of stored data while preserving the reconstruction quality of the JPEG-compressed MSB stream. This design effectively exploits the segment-level redundancy of the structural component and provides an efficient representation for the high-order information of medical images.

### C. Compression of LSB Component

The LSB component mainly contains fine-grained residual variations and local texture details that are complementary to the structural information preserved in the MSB component. Compared with the MSB component, the LSB component generally exhibits weaker spatial correlation and more noise-like intensity fluctuations, making it less suitable for direct compression by conventional hand-crafted codecs. Therefore, we employ a Learned Image Compression (LIC) framework [22] to model these low-correlation and high-detail patterns and obtain a compact representation while preserving the information required for image reconstruction.

The adopted LIC module follows a neural transform coding paradigm with hyperprior and context-based entropy modeling. Specifically, the input LSB component *I*_*L*_ is first mapped into a latent representation *y* through an analysis transform *g*_*a*_(*·*):

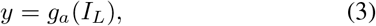

and is then quantized to 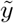. To improve the accuracy of entropy estimation, a hyper-analysis transform *h*_*a*_(·) further extracts a hyper-latent variable *z* from *y*:

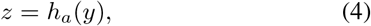

which is quantized to 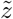 and entropy-coded as an auxiliary bitstream. At the decoder side, 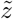 is first recovered and passed through the hyper-synthesis transform *h*_*s*_(·) to generate side information for modeling the distribution of 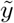.

In addition to the hyperprior information, a context prediction module is used to capture the dependency among previously decoded latent symbols:

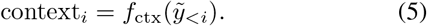

The outputs of the hyperprior branch and the context model are jointly used to estimate the conditional probability distribution of each latent element:

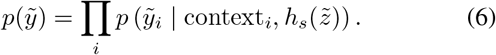

To better characterize the complex distribution of the LSB latent representation, the entropy model employs a Gaussian Mixture Model-based likelihood estimation together with latent-space attention mechanisms [22]. This probabilistic modeling enables more accurate entropy coding than a single-distribution assumption for the irregular residual patterns contained in the LSB component.

Based on the estimated probability distribution, the quantized latent representation 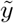is arithmetic-coded to form the main bitstream:

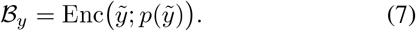

During decoding, the auxiliary bitstream is decoded first to recover 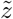, which provides the prior information required for decoding the main bitstream *ℬ*_*y*_. The recovered latent representation 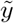 is then passed through a synthesis transform *g*_*s*_(*·*) to reconstruct the LSB component:

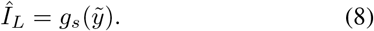

Through the joint use of hyperprior information, contextual dependency modeling, and mixture-based likelihood estimation, the LIC module can efficiently represent the irregular statistical distribution of the LSB component. In this way, the proposed framework preserves low-order residual details with a compact bitstream, complementing the structurally oriented MSB compression branch and enabling accurate reconstruction of the original high bit-depth medical image.

### D. Evaluation Metrics

The performance of the proposed method is evaluated in terms of compression efficiency, and reconstruction fidelity. Compression efficiency is measured by bits per pixel (Bpp) and compression ratio (CR). Since the proposed method generates two compressed streams from the MSB and LSB components, Bpp is calculated as

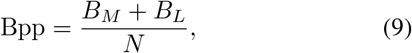

where *B*_*M*_ and *B*_*L*_ denote the number of compressed bits of the MSB and LSB streams, respectively, and *N* is the number of pixels in the original image. The compression ratio is defined as

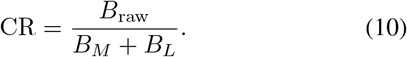

For the 16-bit medical images used in this study, *B*_raw_ = 16*N*. Reconstruction fidelity is evaluated using Peak Signal-to-Noise Ratio (PSNR) and Structural Similarity Index Measure (SSIM) [23]. PSNR is calculated under the 16-bit dynamic range unless otherwise specified.

## IV. Results

### A. Compression performance

We compare the proposed method with two categories of representative compression algorithms: traditional image codecs and learning-based compression models. The traditional group includes JPEG-LS [4], JPEG-2000 [5], PNG [9], JPEG-XL [10], and JP3D [11], while the learning-based group includes L3C [12], LC-FDNet [13], aiWave [14], BCM-Net [15], and BD-LVIC [16].

It should be noted that some of compared methods are numerically lossless or designed for high-fidelity compression, while the proposed method is not strictly pixel-wise lossless. Therefore, the results should be interpreted as a compression-efficiency comparison under reconstruction-fidelity constraints rather than a direct lossless-to-lossless comparison. In addition to Bpp and CR, PSNR and SSIM are reported under the 16-bit dynamic range to verify that the proposed method maintains high pixel-level and structural fidelity after decompression.

As shown in Table II, the proposed method consistently achieves the lowest Bpp and the highest CR across all four datasets. On TRABIT, our method reduces the Bpp to 1.632, compared with 1.856 achieved by the best competing method BD-LVIC. Similar advantages are observed on Covid-CT, Heart-MRI, and Chaos-CT, where our method achieves 2.012, 2.526, and 2.174 Bpp, respectively. Compared with the strongest baseline on each dataset, the proposed method reduces the bit rate by 12.1%, 54.5%, 37.2%, and 52.6% on TRABIT, Covid-CT, Heart-MRI, and Chaos-CT, respectively. In terms of CR, the proposed method achieves 9.809, 7.950, 6.337, and 7.356 on the four datasets, respectively, also outperforming all competing methods.

**TABLE II.** Compression performance comparison on different medical image datasets.

| Method | TRABIT |  | Covid-CT |  | Heart-MRI |  | Chaos-CT |  |
| --- | --- | --- | --- | --- | --- | --- | --- | --- |
|  | Bpp↓ | CR↑ | Bpp↓ | CR↑ | Bpp↓ | CR↑ | Bpp↓ | CR↑ |
| *Traditional codecs* |  |  |  |  |  |  |  |  |
| JPEG-XL | 2.222 | 7.204 | 4.730 | 3.410 | <u>4.019</u> | <u>2.762</u> | <u>4.587</u> | <u>3.556</u> |
| JP3D | 4.177 | 3.831 | 8.789 | 1.826 | 8.381 | 1.901 | 8.488 | 2.065 |
| PNG | 3.083 | 5.190 | 7.090 | 2.257 | 12.034 | 1.330 | 9.012 | 1.775 |
| JPEG-LS | 2.224 | 7.194 | 5.019 | 3.188 | 11.049 | 1.448 | 7.383 | 2.167 |
| JPEG-2000 | 2.579 | 6.204 | 5.318 | 3.009 | 11.436 | 1.399 | 6.872 | 2.328 |
| *Learning-based codecs* |  |  |  |  |  |  |  |  |
| L3C | 2.245 | 7.127 | 5.639 | 2.837 | 10.378 | 1.542 | 6.393 | 2.503 |
| LC-FDNet | 2.122 | 7.540 | 5.395 | 2.966 | 11.124 | 1.438 | 6.018 | 2.659 |
| aiWave | 1.910 | 8.377 | 4.910 | 3.259 | 9.161 | 1.747 | 5.077 | 3.151 |
| BCM-Net | 1.910 | 8.379 | 4.708 | 3.349 | 5.666 | 2.437 | 6.519 | 2.294 |
| BD-LVIC | <u>1.856</u> | <u>8.621</u> | <u>4.421</u> | <u>3.619</u> | 8.842 | 1.810 | 5.012 | 3.192 |
| Ours | **1.632** | **9.809** | **2.012** | **7.950** | **2.526** | **6.337** | **2.174** | **7.356** |

This consistent improvement indicates that explicitly considering the different information characteristics of high-order and low-order bit planes is beneficial for high bit-depth medical image compression. Building on this insight, the proposed hybrid framework leads to more compact representations across different modalities and anatomical regions while maintaining diagnostic fidelity, as further verified in the following subsection.

As shown in Table III, the proposed method achieves high reconstruction fidelity on all datasets. Previous studies have suggested that, under a 16-bit dynamic range, PSNR values of approximately 60–80 dB are commonly observed for lossy image and video compression [24]. In our results, the PSNR values range from 64.53 dB to 75.18 dB, and the SSIM values are all above 0.9996, indicating that the decompressed images maintain high pixel-level and structural fidelity.

**TABLE III.** Quantitative reconstruction fidelity of the decompressed images.

| Dataset | PSNR (dB) | SSIM |
| --- | --- | --- |
| TRABIT | 75.18 | 1.0000 |
| Chaos-CT | 72.32 | 0.9999 |
| Covid-CT | 64.53 | 0.9996 |
| Heart-MRI | 72.76 | 0.9999 |

### B. Model Optimization and Ablation Study

To further analyze the contribution of each component in the proposed framework, we conducted model optimization and ablation experiments on the TRABIT dataset. The experiments focus on three aspects: the selection of the bit-plane decomposition hyperparameter *d*, the compression strategy for the MSB structural stream, and the contribution of each component in the overall architecture. Encoding and decoding times are also reported to provide a reference for computational cost, but the primary objective of this study is to improve storage efficiency while preserving diagnostic fidelity.

#### a) The influence of hyperparameter d

The hyperparameter *d* controls the allocation of information between the MSB and LSB components. A smaller *d* keeps more information in the MSB component, while a larger *d* transfers more low-order information into the LSB component. As shown in Table IV, when *d* = 4, the method achieves higher reconstruction fidelity but produces a higher bit rate, indicating that the MSB stream still contains considerable information to be stored. When *d* = 8, although the bit rate remains relatively low, the reconstruction quality decreases significantly, because excessive information is assigned to the LSB component and then compressed by the lossy learned codec. In contrast, *d* = 6 achieves the lowest Bpp and the highest CR while maintaining a favorable reconstruction quality. Therefore, *d* = 6 is adopted in the final framework as a better trade-off between compression efficiency and reconstruction fidelity.

**TABLE IV.**
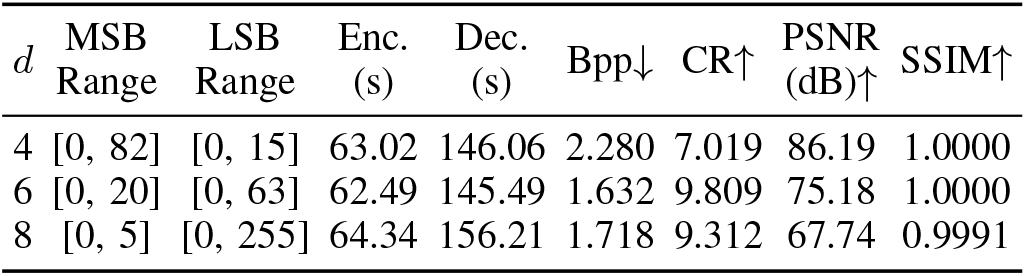
Influence of the bit-plane decomposition hyperparameter *d* on the TRABIT dataset.

| $d$ | MSB Range | LSB Range | Enc. (s) | Dec. (s) | Bpp↓ | CR↑ | PSNR (dB)↑ | SSIM↑ |
| --- | --- | --- | --- | --- | --- | --- | --- | --- |
| 4 | [0, 82] | [0, 15] | 63.02 | 146.06 | 2.280 | 7.019 | 86.19 | 1.0000 |
| 6 | [0, 20] | [0, 63] | 62.49 | 145.49 | 1.632 | 9.809 | 75.18 | 1.0000 |
| 8 | [0, 5] | [0, 255] | 64.34 | 156.21 | 1.718 | 9.312 | 67.74 | 0.9991 |

#### b) Ablation study of the overall architecture

Table V evaluates the contribution of each component in the complete compression framework. Compared with the JPEG-only baseline, adding DSS reduces the Bpp from 5.614 to 5.010 and improves the CR from 2.850 to 3.193, while slightly reducing both encoding and decoding time. This indicates that DSS effectively removes redundant blocks without introducing additional computational burden. After incorporating the LIC-based LSB compression branch, the Bpp is further reduced to 1.632 and the CR increases to 9.809. Although the LIC branch increases the encoding and decoding time, it brings the most significant improvement in compression efficiency. Therefore, the complete architecture achieves a better storage-efficiency trade-off by introducing learned compression only for the LSB component.

**TABLE V.** Ablation study of the overall architecture on the TRABIT dataset.

| Method | Enc. (s) | Dec. (s) | Bpp↓ | CR↑ | PSNR (dB)↑ | SSIM↑ |
| --- | --- | --- | --- | --- | --- | --- |
| JPEG | 8.75 | 9.11 | 5.614 | 2.850 | 76.66 | 1.0000 |
| JPEG+DSS | 8.31 | 8.59 | 5.010 | 3.193 | 76.66 | 1.0000 |
| JPEG+DSS+LIC | 62.49 | 145.49 | 1.632 | 9.809 | 75.18 | 1.0000 |

### C. Application to DNA Storage

Beyond medical image compression, the proposed method can also serve as a front-end compression module for DNA-based medical image storage. Since DNA synthesis cost and storage complexity are closely related to the length of the generated nucleotide sequence, reducing the compressed bitstream before DNA encoding is essential for improving storage efficiency.

For the DNA storage application, the compressed bitstream generated by the proposed image compression framework is further converted into DNA sequences using a dual-rule DNA encoding strategy based on chaotic mapping [25]. This method maps binary data into nucleotide sequences through two encoding rules, while chaotic mapping is used to control the rule selection process and regulate the GC content of the generated sequences. By incorporating such a downstream DNA coding strategy, the compressed bitstream can be converted into DNA sequences with favorable biochemical characteristics.

The DNA storage efficiency is evaluated by the number of nucleotides and the coding density *r*, which is defined as

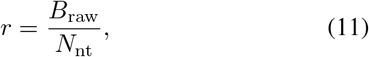

where *B*_raw_ denotes the total number of raw image bits and *N*_nt_ denotes the number of nucleotides required for DNA storage. A larger *r* indicates higher DNA storage efficiency.

Since HELIX [26] and INNSE [27] only support 8-bit inputs, their PSNR and SSIM are evaluated in the 8-bit display domain. DNA-Palette [28] is a fully lossless method and therefore achieves infinite PSNR and an SSIM of 1.0000. For the proposed method, the native 16-bit reconstruction fidelity is reported, consistent with Table III. Therefore, the quality metrics in this table are used as reference indicators, while the main focus of this comparison is DNA storage efficiency.

As shown in Table VI, DNA-Palette achieves exact reconstruction, but requires 24,112,110 nt and yields a relatively low coding density of 4.3133 bits/nt. HELIX-60* achieves the shortest nucleotide sequence and the highest coding density among all methods, but its reconstruction quality is substantially lower, with a PSNR of 28.96 dB and an SSIM of 0.9569. Compared with INNSE, the proposed method reduces the nucleotide number from 13,244,000 to 9,761,499 while improving the PSNR from 41.17 dB to 75.18 dB and the SSIM from 0.9924 to 1.0000. These results indicate that the proposed method provides a favorable balance between nucleotide storage efficiency and reconstruction fidelity.

**TABLE VI.** DNA storage efficiency and reconstruction quality comparison on the TRABIT dataset. * HELIX-60/90/100 represent variants with different JPEG quality factors.

| Method | Number of nt↓ | $r$ (bits/nt)↑ | PSNR (dB)↑ | SSIM↑ |
| --- | --- | --- | --- | --- |
| DNA-Palette | 24,112,110 | 4.3133 | inf | 1.0000 |
| HELIX-60* | 6,661,646 | 15.6122 | 28.96 | 0.9569 |
| HELIX-90 | 12,234,578 | 8.5007 | 27.38 | 0.9583 |
| HELIX-100 | 24,807,400 | 4.1924 | 21.53 | 0.8843 |
| INNSE | 13,244,000 | 7.8528 | 41.17 | 0.9924 |
| Ours | 9,761,499 | 10.6544 | 75.18 | 1.0000 |

Table VII further evaluates the proposed compression framework as an image compression stage before DNA encoding on different medical image datasets. The proposed method achieves coding densities of 10.6544, 8.1129, 9.4999, and 14.2703 bits/nt on TRABIT, Chaos-CT, Covid-CT, and Heart-MRI, respectively. Although the number of generated nucleotides varies with image resolution, slice number, and image content, the compressed bitstreams can be consistently converted into DNA sequences across both MRI and CT datasets. These results indicate that the proposed method is compatible with downstream DNA encoding and has potential for reducing the nucleotide storage burden in DNA-based medical image storage.

**TABLE VII.** DNA storage application results of the proposed method on different medical image datasets.

| Dataset | Enc. (s) | Dec. (s) | Number of nt | $r$ (bits/nt) |
| --- | --- | --- | --- | --- |
| TRABIT (MRI) | 64.09 | 146.84 | 9,761,499 | 10.6544 |
| Chaos-CT | 150.20 | 342.531 | 40,446,351 | 8.1129 |
| Covid-CT | 83.04 | 195.45 | 19,868,067 | 9.4999 |
| Heart-MRI | 90.14 | 174.00 | 22,962,303 | 14.2703 |

Table VIII further evaluates the biochemical characteristics of the DNA sequences generated after applying downstream DNA encoding to the compressed bitstreams of different medical image datasets. Across all datasets, the average GC content remains close to 50%, ranging from 50.25% to 50.41%, and the average melting temperature is highly stable, varying only from 81.50 to 81.57 ^*°*^C. In addition, the maximum homopolymer length is consistently limited to 3 nt. These results show that the bitstreams produced by the proposed compression framework can be smoothly converted by the DNA coding pipeline into sequences satisfying favorable biochemical char-acteristics. This demonstrates the compatibility of the proposed compression method with downstream DNA storage rather than introducing additional constraints or instability during DNA sequence generation.

**TABLE VIII.** Biochemical characteristics of the DNA sequences generated from different medical image datasets.

| Dataset | counts | GC content | Tm (°C) | Max Homo. |
| --- | --- | --- | --- | --- |
| TRABIT (MRI) | 47,157 | 50.37% | 81.55 | 3 |
| Heart-MRI | 110,929 | 50.36% | 81.54 | 3 |
| Covid-CT | 95,981 | 50.25% | 81.50 | 3 |
| Chaos-CT | 195,393 | 50.41% | 81.57 | 3 |

## V. Conclusion

In this paper, we proposed a bit-plane-aware dual-stream compression framework for high bit-depth medical images. By decomposing each 16-bit image into an MSB structural component and an LSB detail component, the proposed method applies JPEG coding with Duplicate Segment Skipping (DSS) to the MSB stream and learned image compression to the LSB stream, thereby exploiting the different statistical characteristics of high-order structural information and low-order residual details. Experiments on four MRI and CT datasets demonstrated that the proposed method consistently achieved superior compression performance over representative traditional and learning-based codecs, while preserving high reconstruction fidelity.

We further evaluated the proposed framework as a preprocessing compression step for DNA-based medical image storage. The results showed that the compressed bitstreams could be effectively integrated with downstream DNA encoding and converted into sequences with stable biochemical characteristics, indicating good compatibility with DNA storage workflows. Overall, the proposed framework provides an effective solution for efficient and diagnostically faithful compression of high bit-depth medical images, with potential application in compact DNA-based medical image archiving.

